# Systematic evaluation of statistical methods for identifying looping interactions in 5C data

**DOI:** 10.1101/201681

**Authors:** Thomas G. Gilgenast, Jennifer E. Phillips-Cremins

## Abstract

Chromosome-Conformation-Capture-Carbon-Copy (5C) is a molecular technology based on proximity ligation that enables high-resolution and high-coverage inquiry of long-range chromatin looping interactions. Computational pipelines for analyzing 5C data involve a series of inter-dependent normalization procedures and statistical methods that markedly influence downstream biological results. A detailed analysis of the trade-offs inherent to all stages of 5C analysis has not been reported, but is essential for understanding the biological basis of looping. Here, we provide a comparative assessment of method performance at each step in the 5C analysis pipeline, including sequencing depth and library complexity correction, bias mitigation, spatial noise reduction, distance-dependent expected and variance estimation, modeling, and loop detection. We present a detailed discussion of methodological advantages/disadvantages at each step and provide a full suite of algorithms, lib5C, to allow investigators to test the range of approaches on their own high-resolution 5C data. Principles learned from our comparative analyses will have broad impact on many other forms of Chromosome-Conformation-Capture-based data, including Hi-C, 4C, and Capture-C.

## Introduction

Higher-order chromatin folding in the three-dimensional nucleus is critically linked to genome function, including transcription (Deng and Blobel, 2014), replication (Rhind and Gilbert, 2013), recombination (Jhunjhunwala et al., 2009), and X chromosome inactivation (Nora et al., 2012). Molecular methodologies based on proximity ligation and deep sequencing, so-called Chromosome-Conformation-Capture (3C) techniques, have revealed that genomes are arranged into a hierarchy of complex configurations (Dixon et al., 2012; Lieberman-Aiden et al., 2009). One unique folding feature is the spatial juxtaposition of linearly-distant genomic loci into long-range contacts termed looping interactions. More than 10,000 looping interactions have been identified genome-wide in the highest-resolution genome-wide 3C study to date in human cell lines (Rao et al., 2014). Efforts are currently underway to map genome-wide looping interactions across many species, cell types and genetic perturbations (Dekker et al., 2017). As genome-wide looping interaction maps become widely available across a range of cell types, the field will transition to perturbation studies required to dissect the organizing principles and mechanistic roles of specific classes of long-range interactions.

The Chromosome-Conformation-Capture-Carbon-Copy (5C) technique is a leading method for mapping long-range looping interactions (Dostie et al., 2006). 5C adds a primer/PCR-based hybrid capture step to the classic 3C procedure to amplify only the ligation junctions of interest across contiguous regions spanning a subset of the genome. The key strength of this technique is high-resolution inquiry of genome contacts across several Mb-sized regions at restriction fragment-level resolution without the high cost of genome-wide Hi-C analysis. Specifically, 5C requires only 10-30 million reads for fragment-level (300 bp – 4000 kb) chromatin contact maps, whereas Hi-C requires up to 6 billion reads to obtain genome-wide maps at a similar resolution (Rao et al., 2014). Thus, 5C has a key utility in allowing researchers to create high-resolution chromatin folding maps at specific genomic region(s) across hundreds of biological conditions and perturbations at a fraction of the cost of Hi-C.

The extent to which looping interactions differ among cell types is currently unknown, in part because the methodologies for identifying long-range interactions in 3C-based sequencing data vary widely across studies and can dramatically influence the results. A systematic comparison of methods for processing, normalizing, and modeling 5C data with the goal of detecting loops has not been conducted. Moreover, no gold-standard set of algorithms for loop detection in 5C data has been reported. Here, we provide a suite of algorithms, lib5C, for direct systematic comparison of multiple methods at each stage in the 5C analysis pipeline, including: (i) sequencing depth and library complexity correction, (ii) bias mitigation, (iii) spatial noise reduction, (iv) distance-dependent expected and variance estimation, and (v) modeling for the goal of loop detection (**Figure S1**). We compare and contrast the strengths and weaknesses of each method and make recommendations for approaches that yield high-confidence looping interactions. Together, our described approaches and freely available lib5C tools allow for the sensitive and specific quantitative detection of bona fide looping signal in a given biological condition.

## Results

Proximity ligation techniques (e.g. Hi-C, 5C, 4C, CaptureC) capture many types of non-biological signal along with bona fide looping interactions (Yaffe and Tanay, 2011). Biases vary in their type and severity depending on the method and require modeling and correction prior to biological interpretation. In the specific case of 5C, possible artifacts or confounding signal include: (1) sequencing depth and library complexity differences due to technical artifacts and/or batch effects (**Figures 1A-1C**), (2) biases caused by the intrinsic properties of the restriction fragments queried by the assay (including their length and GC content) (**Figures 1D-1F**), (3) spatial noise due to 5C primer design and library complexity (**Figures 1G-1H**), and (4) the expected background signal at each length scale, which varies as a function of spatial genomic distance and region-specific topologically associating domain (TAD) and subTAD structure (**Figures 1I-1K**). Upon correction of these features, it is also critical to understand the mean-variance relationship and parameterize the appropriate statistical model to assign p-values to each possible interaction (**Figures 1L-1N**). Finally, bona fide looping interactions are called as clusters of highly significant pixels and explored for downstream enrichment of epigenetic marks on the linear 10 nm chromatin fiber (**Figures 1O-1Q**). By rigorously modeling the data, investigators can distinguish 3D chromatin loops from other genome folding patterns as well as background and technical noise, thus providing the opportunity for discovery of biological mechanisms governing looping.

**Figure 1:**
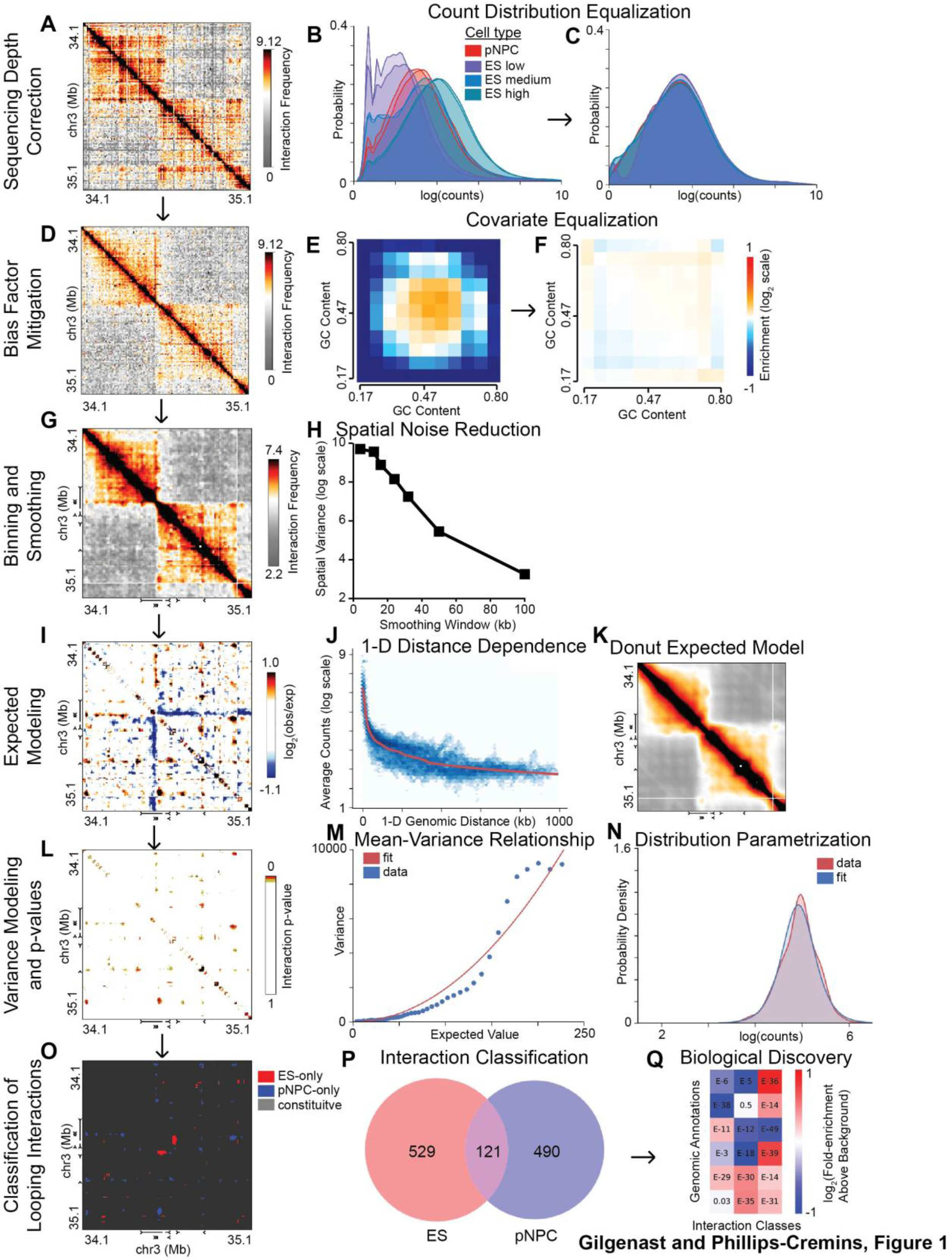
Schematic of 5C analysis pipeline. Flowchart illustrating key steps in 5C analysis. (A) Interaction frequency heatmap of the Sox2 region in primary neural progenitor cells (pNPCs) after sequencing depth correction. (B) 5C count distributions before sequencing depth correction. (C) 5C count distributions after sequencing depth correction. (D) Interaction frequency heatmap of the Sox2 region in pNPCs after bias factor mitigation. (E) GC content bias profile before bias mitigation. (F) GC content bias profile after bias mitigation. (G) Interaction frequency heatmap of the Sox2 region in pNPCs after binning and smoothing. (H) Graph showing spatial variance of the binned contact matrix as a function of the width of the smoothing window used during binning. (I) Heatmap of the Sox2 region in pNPCs showing enrichment (red) and depletion (blue) of contacts relative to an expected model. (J) Illustration of a one-dimensional distance dependence model, which describes the average interaction frequency as a function of linear genomic separation. (K) Expected interaction frequency heatmap for the Sox2 region in pNPCs after distance dependence modeling and donut correction. (L) Heatmap of the Sox2 region in pNPCs showing interaction p-values. (M) Graph showing relationship between expected contact frequency and variance of contact frequency. (N) A log-logistic distribution parameterized using the mean-variance relationship at an expected value of 150 (blue) overlayed with a kernel density estimate of the distribution of real data points with similar expected values. (O) Heatmap of the Sox2 region in pNPCs showing classified interactions. (P) Venn diagram showing the numbers of ES-specific, pNPC-specific, and constitutive interactions. (Q) Enrichment heatmap showing relationships between classes of significant interactions (rows) and various genomic annotations (columns).

Like all genomics assays (Daley and Smith, 2013; Marinov et al., 2014; Sims et al., 2014), 5C libraries can exhibit large differences in complexity and sequencing depth due to technical variation in ligation and fixation efficiency among experimenters, reagents, and protocols. We observed that technical 5C replicates from the same biological condition can show a high degree of variability in their raw counts distribution (Raw, **Figure 2A**), distance-dependent expected counts distribution (Raw, **Figure 2B**), spatial complexity (Raw, **Figure 2C**), and relationship between the raw interaction counts and guanine-cytosine (GC) content of the fragments hybridizing to 5C primers (Raw, **Figure 2D**). Differences in count distributions, distance-dependent expected curves, and spatial complexity trend more with technical batch than with biological conditions (Raw, **Figures 2A-2C**), suggesting that they are driven by artifacts rather than biologically-driven effects. Thus, raw 5C counts exhibit biases caused by library complexity and sequencing depth that require correction prior to biological interpretation.

**Figure 2:**
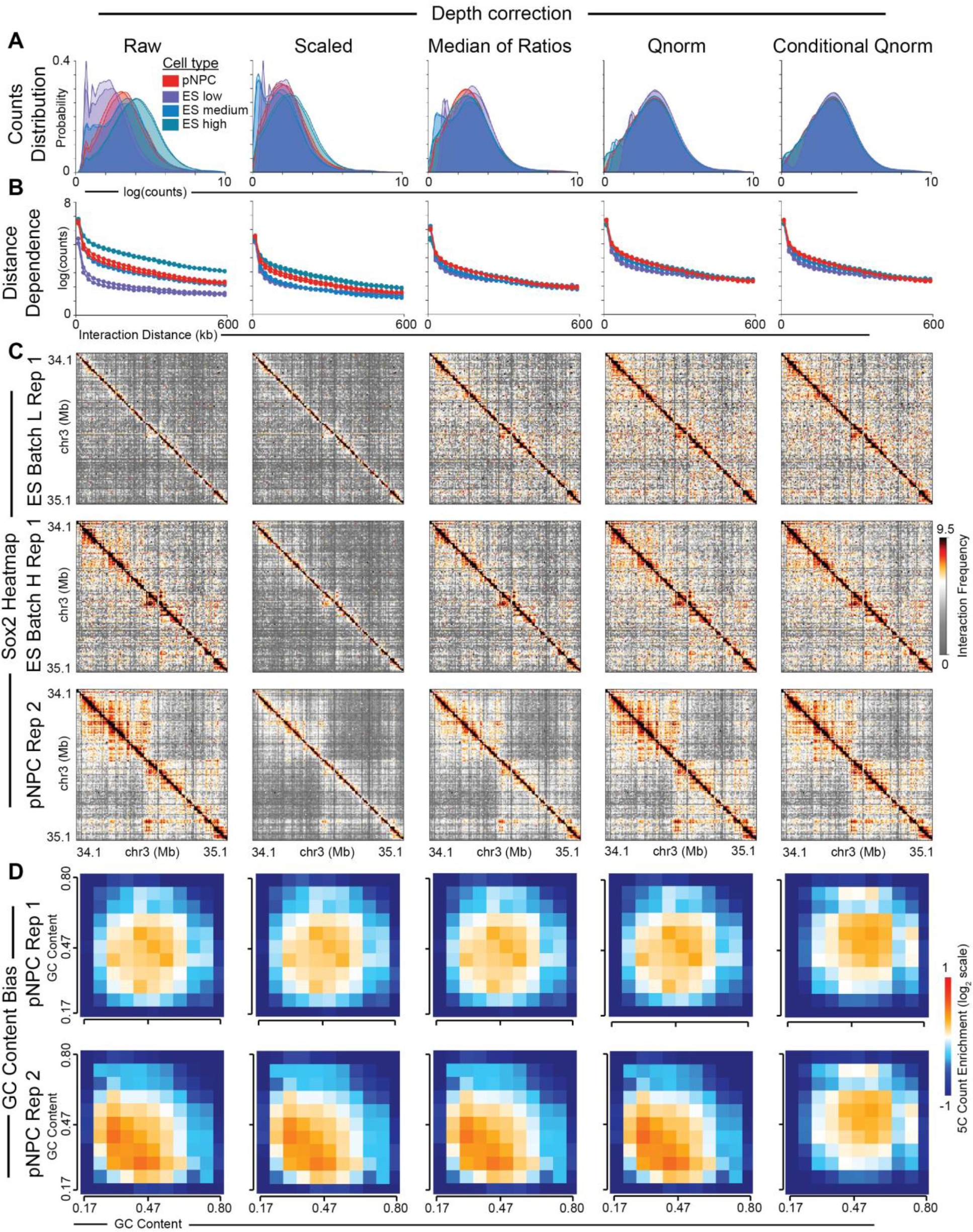
5C sequencing depth correction methods. (A) Count distributions across replicates from various cell types before and after application of sequencing depth correction procedures. Replicates from pNPCs are shown in red while replicates from ES cells are shown in various shades of blue, according to relative sequencing depth (low, medium, or high). (B) Similar comparison for distance dependence curves. (C) Fragment-level contact frequency heatmaps of the region around the Sox2 gene. (D) Heatmaps showing GC content bias profiles of selected replicates from the same cell type. The color indicates the average relative enrichment of ligation detection events as a function of the GC content of the primer designed to one of the participating fragments (x-axis) and that of the other fragment (y-axis).

To compare looping interactions between biological conditions, it is essential to correct for differences in library complexity and sequencing depth. We find that correcting raw counts by a scalar value of total sequencing reads is unable to correct for replicate differences in the shape of the raw counts distribution and distance-dependent expected curve (Scaled, **Figures 2A-2C**). By contrast, correction via the median-of-ratios scaling technique (Anders and Huber, 2010) or quantile normalization can more rigorously normalize the raw counts distributions and distance-dependent expected curves (Median-of-Ratios and Qnorm, **Figures 2A-2C**). We find that equalizing distributions among replicates of the same condition does not sufficiently equalize GC content bias profiles. Indeed, the relationship between raw interaction signal and the GC content of the ligation junctions remains widely variable between replicates even after quantile normalization and median-of-ratios correction (**Figure 2D**). These data indicate that most published methods for correcting for sequencing depth and library complexity differences in proximity ligation data are insufficient to account for intra-condition technical variation among replicates.

To account for the strong replicate-specific effect of GC content bias on raw interaction count, we developed and applied a variation of the conditional quantile normalization method proposed by Hansen and colleagues for RNA-seq (Hansen et al., 2012). Specifically, we stratified all pairs of restriction fragments by the GC content of the portion of the DNA sequence homologous to the 5C primers. We conditionally quantile normalized ligation junctions in each GC content strata conditionally across all 5C replicates from all biological conditions. The conditional quantile normalization procedure fully corrected 5C libraries for replicate-specific distributional differences (conditional qnorm, **Figures 2A-2C**) and GC content bias profiles (conditional qnorm, **Figure 2D**) without any distortion to the underlying heatmaps (conditional qnorm, **Figure 2C**). Altogether, our new conditional quantile normalization method offers robust correction for sequencing depth and library complexity differences between technical replicates without negatively affecting the underlying condition-specific genome folding patterns.

Although GC content profiles have been equalized between samples after conditional quantile normalization (**Figure 2D**), each individual sample still exhibits strong fragment-dependent GC content biases that must then be corrected prior to the detection of looping interactions (**Figure 3**). Hi-C ligation junctions are also known to exhibit read count biases linked to the intrinsic properties of the fragments (e.g. GC content, fragment length, and mappability) (Jin et al., 2013; Yaffe and Tanay, 2011). The effects of intrinsic biases are not localized to particular pairs of interacting fragments, such as those engaged in looping interactions, but instead increase or decrease the raw interaction counts for all ligation partners of the fragment in question. Thus, intrinsic biases are made manifest as “lines” of under-or over-enriched counts spanning a significant proportion of the raw fragment-fragment contact matrices. Visual inspection confirmed that bias ‘lines’ also exist in 5C data (Raw, **Figure 3A**). We also observed this phenomenon more quantitatively as a wide dynamic range (over 80-fold difference in medians) of interaction count profiles among the restriction fragments (Raw, **Figure S2A**). An important consequence of fragment bias is that it can obscure biological signal due to looping events, as evidenced by zoom-in heatmaps of two bona fide looping interactions previously reported in (Beagan et al., 2016) (Raw, **Figure 3B**). Together, these data highlight that intrinsic fragment biases should be corrected before calling significant biological interactions in 5C data.

**Figure 3:**
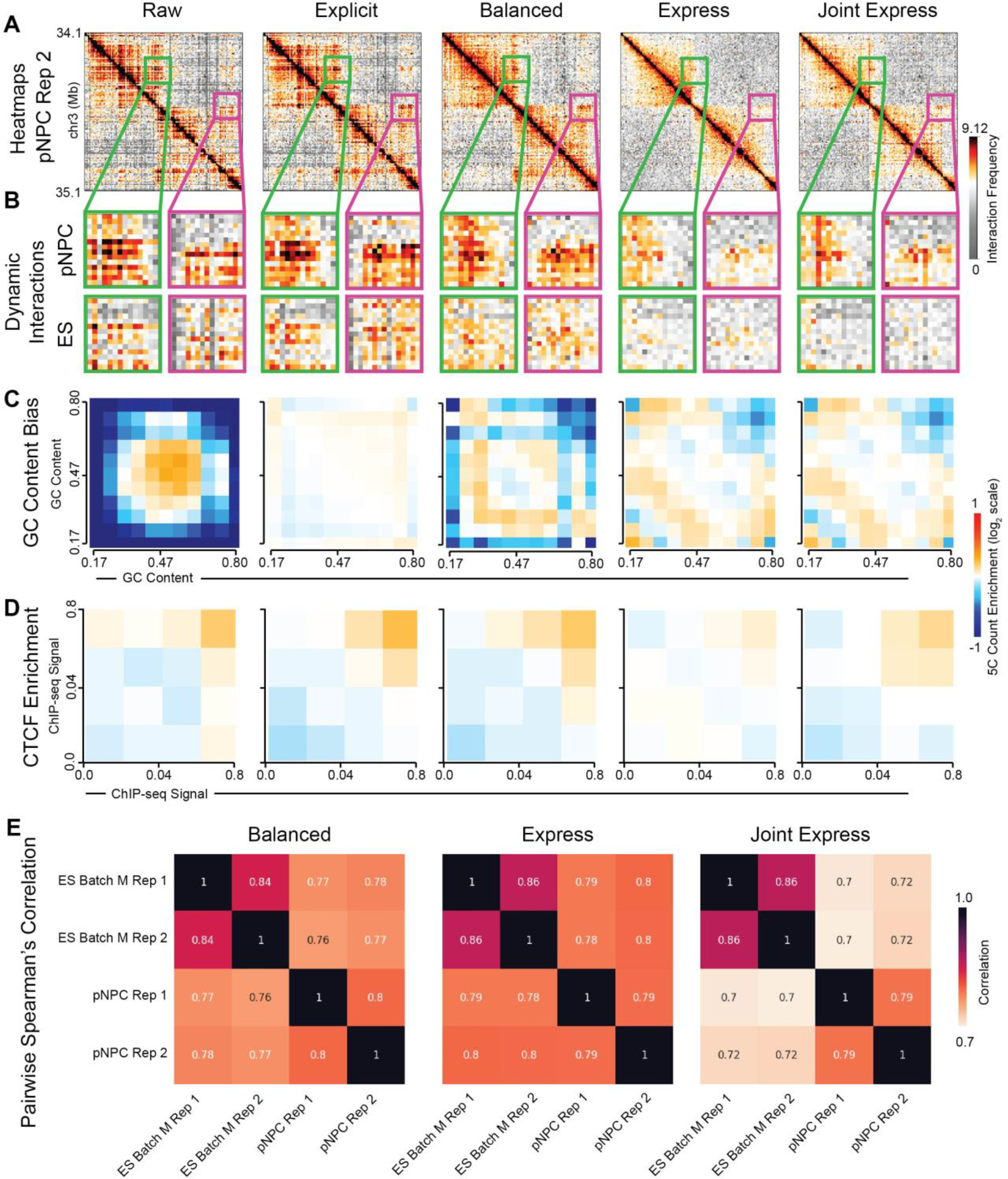
5C bias factor mitigation procedures. (A) Fragment-level contact frequency heatmaps of the region around the Sox2 gene in pNPCs. (B) Zoom-in views around previously identified pNPC-specific interactions between the Sox2 gene and NPC-specific enhancers. The upper row shows the interaction profile in pNPCs, while the lower row shows the same window in ES cells. (C) Heatmaps showing GC content bias profiles before and after bias factor mitigation. The color indicates the average relative enrichment of ligation detection events as a function of the GC content of the primer designed to one of the participating fragments (x-axis) and that of the other fragment (y-axis). (D) Similar to (C), but stratifying fragment-fragment ligation junctions according to CTCF ChIP-Seq signal over the fragment (computed as the average number of reads per million mapping to a 4 kb window centered at the fragment midpoint). (E) Tables showing pairwise Spearman’s correlation coefficients between replicates after ICED matrix balancing (left), after Express normalization (center), and after Joint Express normalization (right).

Many approaches to correcting intrinsic fragment artifacts in Hi-C data have been reported, but the performance of the correction methods (i.e., explicit bias factor modeling, Knight-Ruiz or ICED matrix balancing, and Express matrix balancing) has not been systematically assessed in 5C data. We first explored our 5C data for possible previously reported Hi-C biases due to primer GC content and fragment length (Jin et al., 2013; Yaffe and Tanay, 2011). Consistent with previous reports, we observed a strong under-enrichment for detected ligation junctions between fragments with extreme GC content (Raw, **Figure 3C**). We also observed a trend toward higher detection of ligation junctions between larger fragments (Raw, **Figure S2B**). Finally, we stratified fragments according to the normalized ChIP-seq signal in a 4 kb interval centered on the fragment midpoint and found that ligation junctions tend to exhibit stronger interaction frequency when CTCF signal is high in both fragments (Raw, **Figure 3D**). These results are consistent with previous findings that CTCF anchors the base of looping interactions genome-wide (Beagan et al., 2017; Li et al., 2010; Rao et al., 2014; Sanyal et al., 2012; Tang et al., 2015). Thus, both candidate bias factors and bona fide epigenetic marks are covariates which may contribute to the 5C interaction counts.

We next surveyed a variety of methods for attenuating GC content and fragment length biases while keeping the known biological link between CTCF and interaction strength intact. We first explicitly modeled and corrected the conditional quantile normalized counts for GC content and restriction fragment length biases (detailed in **STAR Methods**). This approach is conceptually similar to seminal works in which biases were explicitly modeled and corrected in Hi-C data (Jin et al., 2013; Yaffe and Tanay, 2011). Although GC content and restriction fragment length bias factors were almost completely attenuated after correction (Explicit, Figures 3B **and S2B**), the ‘lines’ in the heatmaps remained apparent and fragments still showed a wide dynamic range of counts (Explicit, Figures 3A **and S2A**). The core assumptions behind explicit modeling approaches are that (i) all the intrinsic factors contributing to technical bias are known and (ii) their influence on detected ligation junction counts can be modeled reasonably well with tractable functions. These results suggest that there are still unknown bias factors in 5C data that are not captured by GC-content and restriction fragment length correction.

We next compared explicit modeling to matrix balancing algorithms that implicitly correct for biases without directly defining their specific sources (Imakaev et al., 2012; Knight and Ruiz, 2013; Rao et al., 2014; Sauria et al., 2015). Matrix balancing algorithms have been effectively applied to Hi-C data (Crane et al., 2015; Imakaev et al., 2012; Rao et al., 2014), and depend on the assumptions that (i) all fragments throughout the genome have “equal visibility” (i.e. equal propensity for detection via a proximity ligation assay) and (ii) the intrinsic fragment-specific biases can be represented as a single scalar value for each fragment that interacts multiplicatively with the intrinsic biases of its ligation partners. An open unanswered question is whether these assumptions apply to 5C data given that the genomic regions are relatively small (1-10 Mb) and biases may follow nonlinear relationships. We first applied ICED matrix balancing to the conditional quantile normalized 5C counts. We observed that lines in the heatmaps are strongly attenuated while preserving the cell-type specific looping interactions (Balanced, Figures 3A-3B **and S2A**). Importantly, the smooth heatmaps were achieved despite no effect on the residual fragment length bias and only partial reduction in GC content bias, indicating that biases could be coming from other unknown sources (Balanced, Figures 3C **and S2B**). We also investigated the Express matrix balancing algorithm, which additionally incorporates information from a distance-dependent expected model in the computation of the bias factors (Sauria et al., 2015). We find that Express provides the strongest ubiquitous correction of GC content bias, fragment length bias and lines in heatmaps, but was too harsh in attenuating real biological looping interactions (Express, Figures 3A-3C **and S2A-S2B**). Consistent with these results, Express resulted in marked attenuation of the known enrichment of CTCF at the base of the strongest interacting fragment-fragment ligation junctions (**Figure 3D**). Together these data indicate that Express matrix balancing can lead to over-normalization of 5C data from at least some alternating primer designs and loss of meaningful biological looping interactions.

We created a variant called Joint Express to account for Express’ over-smoothing while retaining its ability to attenuate bias factors. The canonical Express algorithm computes a unique bias vector for each replicate. Our new Joint Express variant computes a single bias vector by integrating information from all replicates (see **STAR Methods**). It relies on the assumptions that (i) variation in technical bias is negligible among the replicates and (ii) any observed bias vector differences correspond to biologically meaningful differences in cell type-specific interaction frequency. We see that Joint Express preserves known biological looping interactions and mostly smooths lines in heatmaps similar to ICED (Joint Express, Figures 3A-3B **and S2A**), while offering small improvements in the smoothing of GC content and fragment length biases compared to ICED (Joint Express, Figures 3C **and S2B**). We also observe that Joint Express improves CTCF enrichment at high interaction frequency ligation junctions compared to Express (**Figure 3D**). Finally, counts processed using the Joint Express algorithm retained a greater degree of cell type specificity as evidenced by weaker inter-replicate correlations than those processed by canonical Express or ICED (**Figure 3E**). Altogether, these data indicate that traditional matrix balancing by ICED or its variants and our new variant on the Express algorithm, Joint Express, represent the highest performing methods for removing technical fragment-specific biases from 5C data while preserving biologically-important interactions.

Hi-C data is typically binned using non-overlapping windows of a predetermined width and summing the detected ligation events within each window. An element of such a binned matrix can be readily interpreted as the total number of times a ligation junction was detected between the two bins in a population of cells, regardless of which specific restriction fragments were involved in the detected ligation. 5C data is particularly susceptible to spatial noise due to the alternating nature of most primer designs which does not comprehensively query all possible ligation junctions. Thus, instead of using non-overlapping bins tiling the genomic region of interest, we employ a sliding window where the “step” between successive evaluations of the window’s interaction frequency is smaller than the window width (**Figure 4**). Region-wide heatmaps binned with a small smoothing window (4 kb, corresponding to no overlap between neighboring windows) show a large degree of spatial heterogeneity, including many interspersed regions where no data are available (**Figure 4A**). By visually inspecting zoomed-in heatmaps, we found discontinuous and noisy interaction signal at a known looping interaction connecting the Sox2 gene with a putative NPC-specific enhancer marked by NPC-specific H3K27ac (**Figure 4B**). We increased the smoothing window to 16 kb with a 4 kb step and observed markedly reduced spatial noise and clear emergence of punctate loops. Noteworthy, very large smoothing windows markedly attenuate spatial noise and highlight TAD/subTAD structure, but at the expense of severe over-smoothing and complete loss of looping interactions (**Figure S3B**). Consistent with these findings, increasing the smoothing window size results in a decreased number of bin-bin pairs with strong interactions (**Figure 4C**) and an inverse relationship with the spatial variance, or noise, observed in the binned contact matrix (Figure 4D, **STAR Methods**). Together, these data suggest that by adjusting the size of the binning window, the investigator can strike a balance between spatial noise in the contact matrices and the ability to detect features at high resolution.

**Figure 4:**
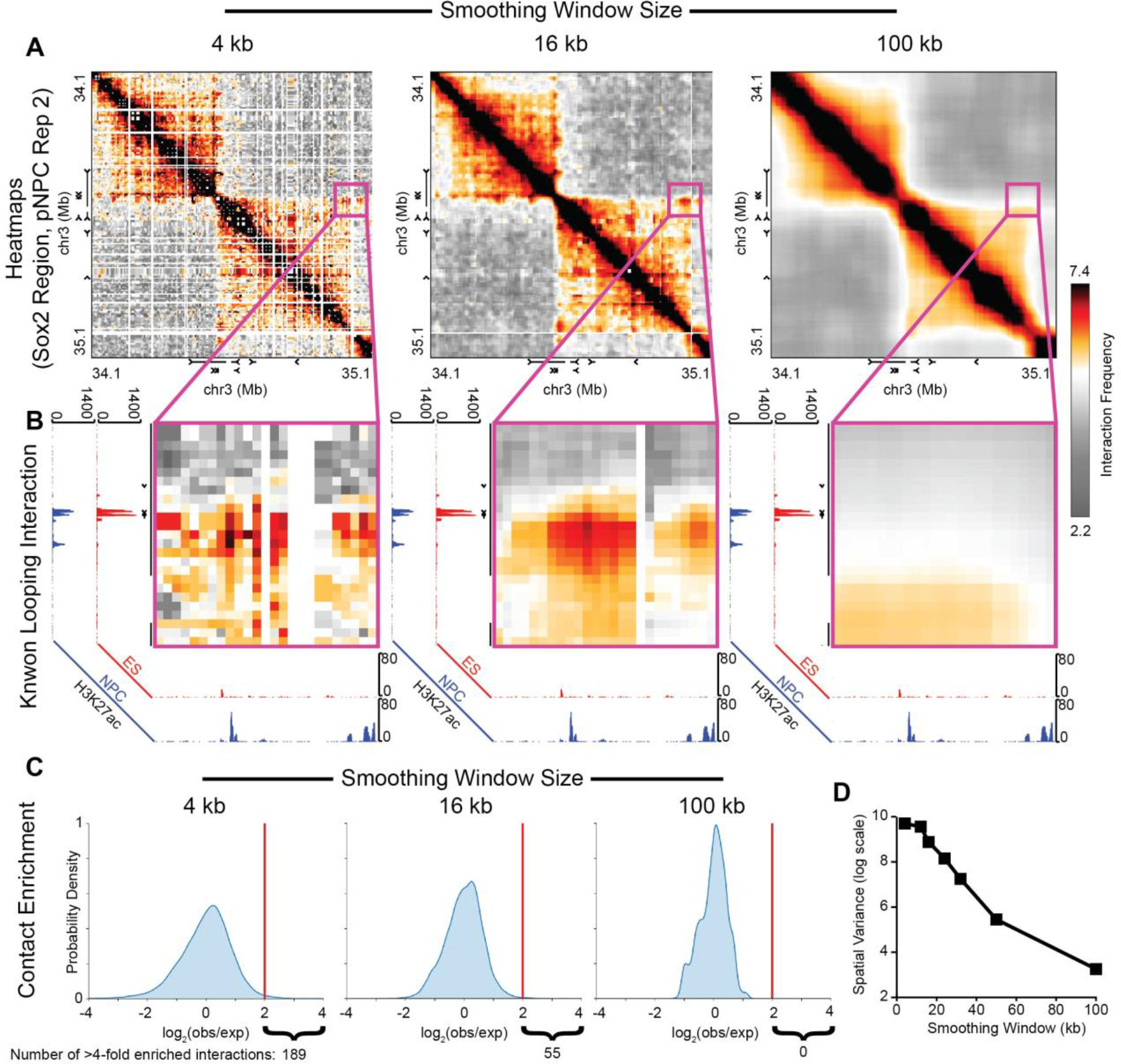
Effects of binning and smoothing on 5C contact matrices. (A) Contact frequency heatmaps of the Sox2 region in pNPCs, binned at 4 kb matrix resolution with a variety of smoothing window widths (from left to right: 4 kb, 16 kb, 100 kb). (B) Zoom-in view around previously identified interaction between the Sox2 gene and an NPC-specific enhancer. ChIP-seq tracks show H3K27ac signal in ES cells and NPCs. (C) Distributions of the enrichment of smoothed values over a simple distance-dependent expected across the Sox2 region. The curly braces indicate the number of bin-bin pairs with smoothed values greater than four times the expected at their interaction distance. (D) Spatial variance of the contact matrix plotted as a function of smoothing window size.

The binned interaction counts (now corrected for sequencing depth, library complexity, intrinsic fragment-specific biases, and spatial noise) exhibit a genomic distance-dependent interaction signal (Lieberman-Aiden et al., 2009) that must be correctly modeled before detecting loops (Rao et al., 2014). This so-called distance-dependent background is made manifest as a strong band of high interaction counts along the diagonal of genome folding heatmaps (**Figure 5A**). We first applied a simple log-log regression to model the changes in expected interaction value between two loci as a function of the linear genomic distance (Figure 5B, **STAR Methods**). Alternative modeling options provided very similar fits (**Figure S4A, STAR Methods**). A critical decision in modeling the distance-dependence expectation is whether a single, global fit to all the data or a 5C region-specific fit should be applied. We empirically observed that different 5C regions exhibit dramatically different distance dependence relationships (**Figure S4B**). We found a specific example of this in the 5C region around the Nanog locus, where the global expected model overestimates the average counts at every distance scale (**Figure 5B**). Thus, the regional outperforms the global distance-dependence expected model for correcting the diagonal prior to loop calling.

**Figure 5:**
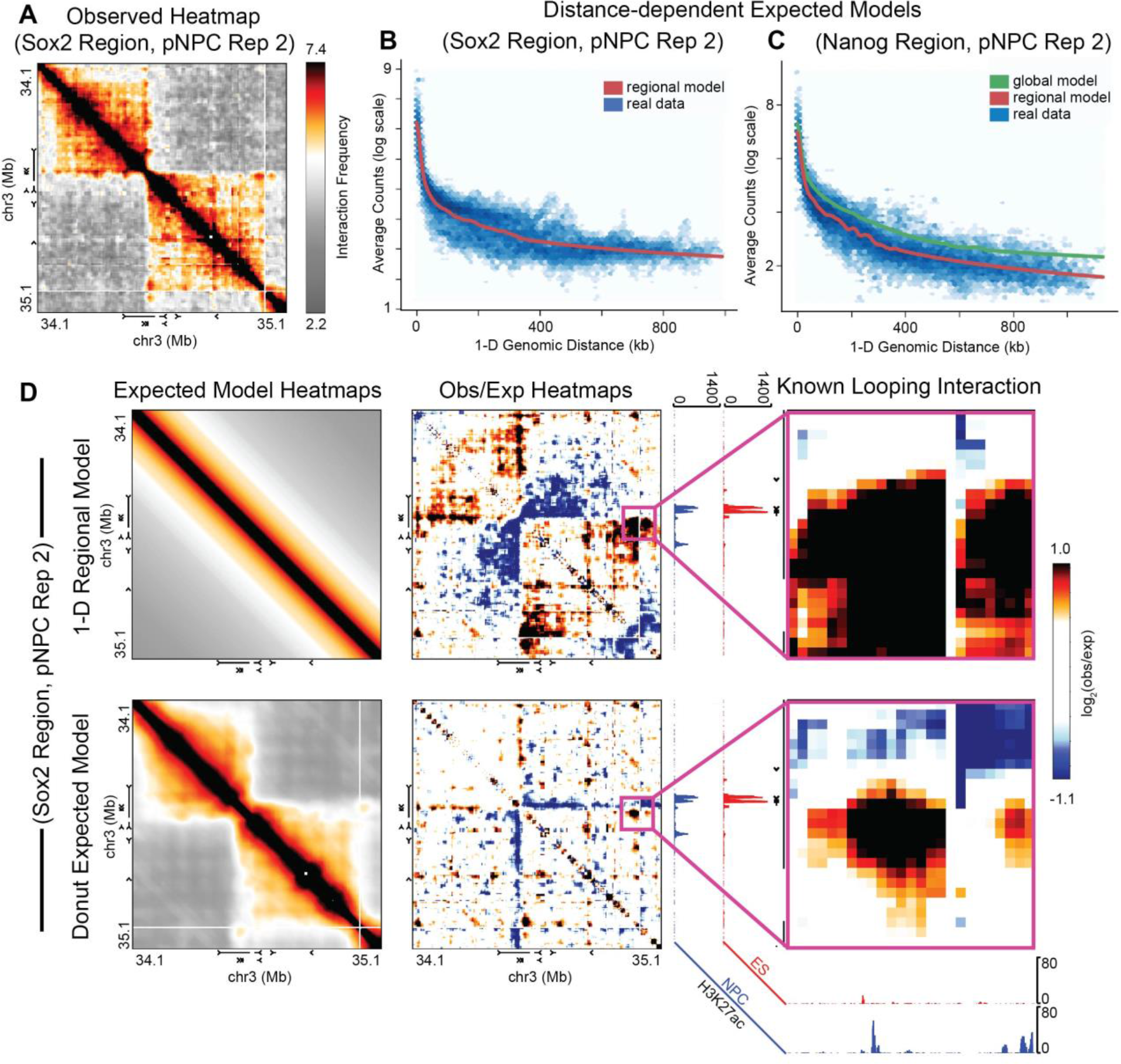
Strategies for modeling the expected 5C counts at each genomic distance scale. (A) Illustration of expected modeling procedure. The entries in the smoothed contact matrix (pixels in the contact frequency heatmap) are shown in blue hexagonal bins on the right. Distance-dependent expected models attempt to fit a function (red curve) through the data (blue points). (B) The plot on the left shows the smoothed contact matrix entries from the Nanog region (blue points) compared to two different distance-dependent expected models: one model fitted only to the data from the Nanog region (red curve) and one fitted to data from all 5C regions in the dataset (green curve). The heatmaps on the right show fold enrichments of the smoothed contact matrix entries over the two distance-dependent expected models. (C) The rows compare a simple one-dimensional distance-dependent expected model (top row) to a donut expected model (bottom row). The left column shows a contact frequency heatmap visualization of the two models at the Sox2 region. The center column shows fold enrichments of the smoothed contact matrix entries over the expected models. The right column shows a zoom-in view around a previously identified interaction between the Sox2 gene and an NPC-specific enhancer. ChIP-seq tracks show H3K27ac signal in ES cells and NPCs.

In addition to distance dependence, the TAD/subTAD architecture strongly influences contact frequencies observed in 5C datasets. Two loci in the same TAD/subTAD tend to interact more frequently than a pair of loci which span a domain boundary, even when these pairs of loci are separated by the same linear genomic distance (Dixon et al., 2012). It is essential to include TAD/subTAD structure in the expectation when detecting long-range looping interactions. To model the expected counts due to chromatin domains, we applied the donut expected filter proposed by Aiden and colleagues (Rao et al., 2014). The donut filter provides a model of the distance-dependent background and local domain structure for each region without having to know the location of TADs/subTADs a priori (**Figure S4C, STAR Methods**). Correcting binned data for the donut expected results in the clear elucidation of punctate loops, whereas relying exclusively on a one-dimensional distance model tends to lead to smearing of punctate looping pixels, especially across the corners of TADs/subTADs (**Figure 5C**). Overall, we see that the donut expected offers the most rigorous and accurate expected model for calling looping interactions in 5C data.

We next investigated methods for estimating the variance under the null model that any given pixel is not engaged in a bona fide looping interaction. More specifically, our null hypothesis is that the observed interaction value at a given pixel is not significantly higher than the expected value at that pixel, when judged according to an estimated variance. The most widely used loop calling method (Rao et al., 2014) models counts with the Poisson distribution under the assumption that the mean is equal to the variance (first panel, **Figure 6A**). We first used the local donut window to directly estimate a sample variance independently for each pixel. We observed that the local variance estimate tended to increase sharply with increasing expected value, however the spread of the data points around this trend was too wide to obtain a robust estimate of the mean-variance relationship (second panel, **Figure 6A**). We next computed the sample variance of groups of pixels with similar expected values within each region, an approach reminiscent of the variance modeling performed in a previously-published Hi-C analysis (Jin et al., 2013). We observed that the trend could consistently be fitted with a simple quadratic function relating the mean and the variance (**Figure S5A**). We therefore fitted one quadratic mean-variance relationship in this fashion for each replicate across all regions (third panel, **Figure 6A**). Although many other approaches could be pursued to model the 5C mean-variance relationship, it is beyond the scope of this paper to pursue additional options.

**Figure 6:**
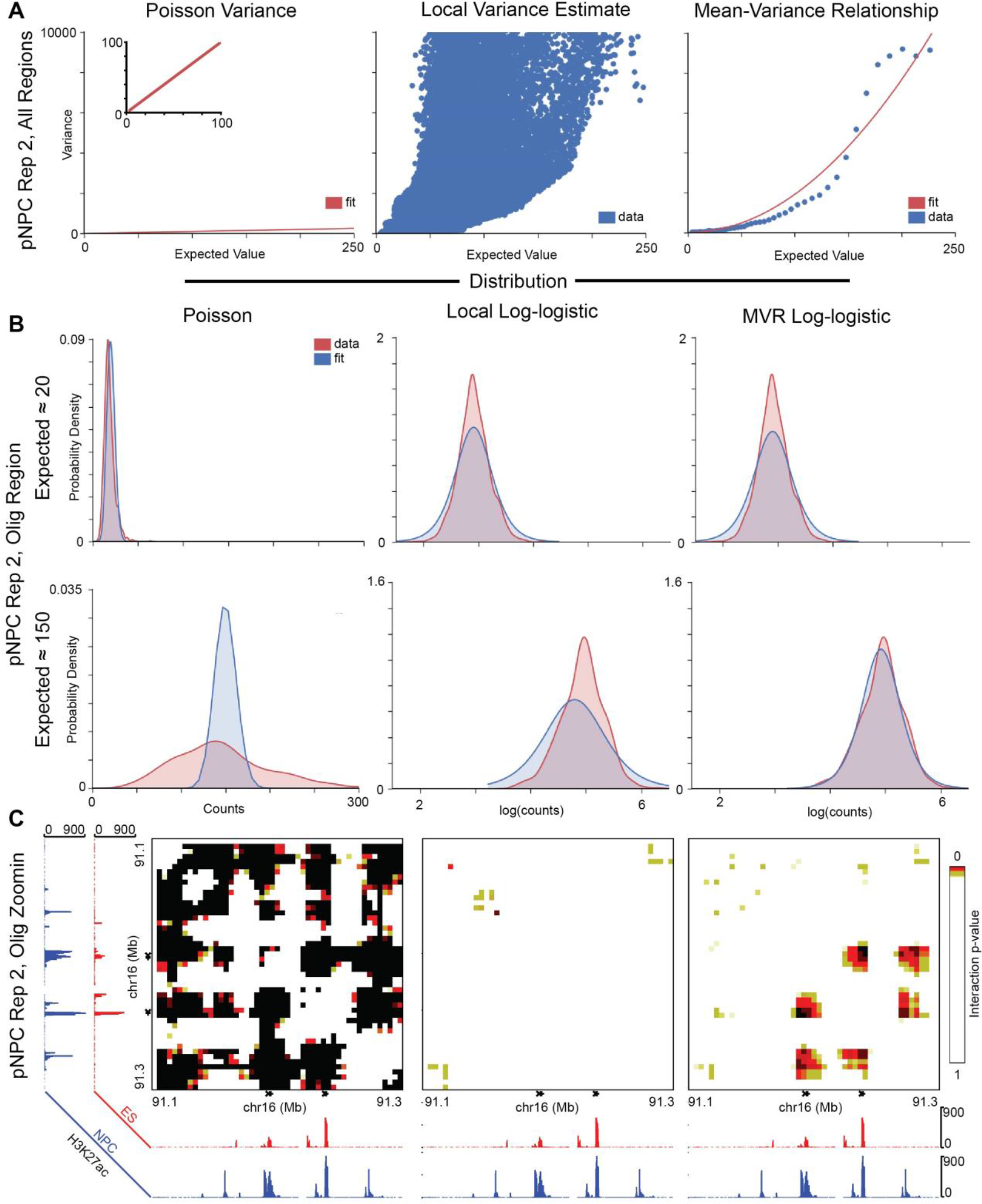
Strategies for modeling the mean-variance relationship in 5C data. (A) From left to right: the Poisson variance model (under which the variance always equals the expected value), a local variance model (under which the variance is estimated using nearby entries in the contact matrix, irrespective of the expected value), and a mean-variance relationship model (under which the variance is assumed to be a quadratic function of the expected value, fitted to the sample variance of contact matrix entries with similar expected values. (B) Theoretical distributions (blue) parametrized according to the variance models in (A) compared to the distribution (red) of contact matrix entry smoothed values with expected values near 20 (top row) and near 150 (bottom row). (C) Heatmaps (zoomed in on the Olig1 and Olig2 genes) of interaction p-values derived from the three distribution parametrization options (from left to right: Poisson, local log-logistic, and MVR log-logistic). P-values are computed against the null hypothesis that the interaction’s value is not higher than expected given the estimated variance.

To assess the performance of the variance estimates, we plotted theoretical distributions parametrized with a specific pixel’s expected value and estimated variance against the distribution of the observed data for pixels with similar expected values **(Figure 6B)**. We observed that the Poisson variance model worked reasonably well for pixels in our 5C dataset with relatively low expected values (e.g., approximately 20, Figure 6B, **upper left**), but seemed to underestimate the variance at higher expected values (e.g., around 150, Figure 6B, **lower left**). We also observed that the local variance estimate can result in over-or under-estimating the variance (e.g., the expected value around 150, Figure 6B, **middle**). Our quadratic MVR model yielded parametrized distributions which more consistently resembled the distribution of the data (Figure 6B, **right**). We also computed p-values based on the parametrized distributions for each pixel, and visualized these as p-value heatmaps **(Figure 6C)**. We observed similar trends as **Figure 6A-B**, with the Poisson model overestimating pixel significance, and the local variance estimate underestimating pixel significance compared to the MVR approach. The quadratic mean-variance relationship assumption leveraged here is conceptually reminiscent of the log-transform performed on observed and expected counts values in earlier 5C analyses (Beagan et al., 2017; Beagan et al., 2016), which can be re-interpreted as a variance stabilizing transform and results in similar p-values (**Figures S5B-S5D, STAR methods**). We also briefly assessed the influence of the choice of the statistical distribution parametrized with a particular mean and variance estimate. We observed that both discrete (such as the negative binomial) and continuous (such as the log-normal) distributions reflected the distribution of the observed data (**Figures S5B-S5C**). For our final analysis, we selected the quadratic MVR approach with parametrized log-logistic distributions.

Finally, we categorized and clustered significant looping interactions and assessed the quality and biological relevance of the called loops. We first applied a threshold at a p-value of 0.05 to decide the significance of individual pixels in individual replicates (**Figure 7A**). We then discarded interactions which were found to be significant in only one of the two replicates for each condition. Next, we classified interactions as ES-only, pNPC-only, or constitutive depending on which conditions they were found to be significant in. Finally, we clustered the classified interactions, since multiple adjacent significant pixels often represent one underlying biological interaction (**Figure 7B**, additional loci highlighted in **Figures S6A-S6D**). All together, we identified 529 ES-specific, 490 pNPC-specific, and 121 constitutive interactions (**Figure 7C**) which were distributed across the 5C regions (**Figure 7D**). Classified interactions showed reasonable enrichments when compared to one-dimensional genomic annotations. For example, ES-specific CTCF and enhancers were enriched in ES-specific loops, constitutive and pNPC-specific loops were enriched for constitutive CTCF, and pNPC-specific YY1 and enhancers were enriched in pNPC-specific loops (**Figure 7E**), consistent with previously published observations (Beagan et al., 2017; Beagan et al., 2016). Finally, we assessed the enrichment of occupied CTCF motifs with different orientations under pNPC-specific loops (**Figure 7F**, ES-specific loops in **Figure S6E**) and observed a strong enrichment for convergently-oriented CTCF motifs, as previously reported (Rao et al., 2014). Thus, our selected analysis conditions at each stage in the 5C pipeline resulted in strong enrichment for occupancy of well-characterized epigenetic marks and transcription factors.

**Figure 7:**
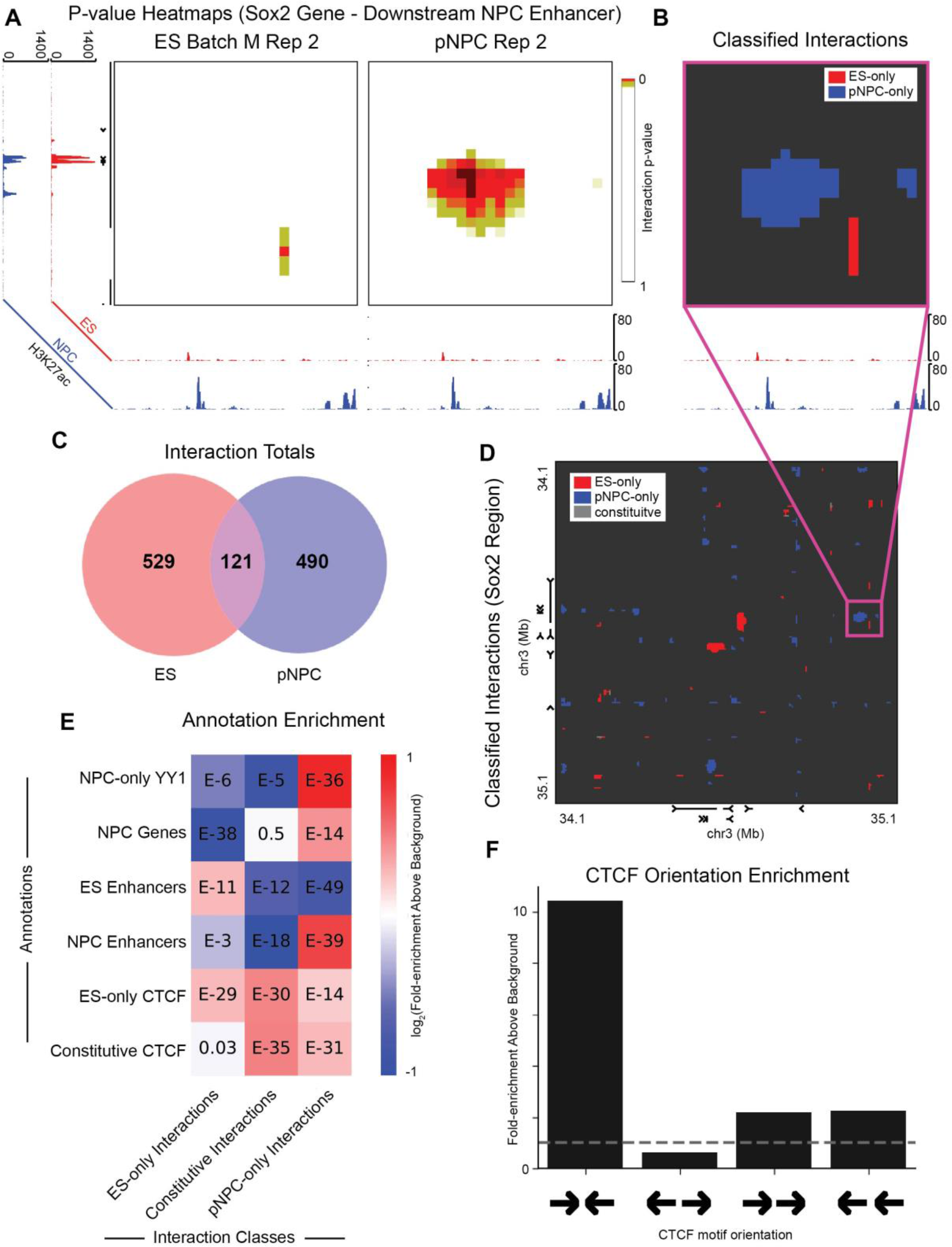
Characterization of statistically significant interactions for enrichment of known epigenetic marks. (A) Zoom-in view around a previously identified interaction between the Sox2 gene and an NPC-specific enhancer. Colors correspond to the significance of the interaction (right tail p-value). ChIP-seq tracks show H3K27ac signal in ES cells and NPCs. (B) Same view as in (A), but colored according to cell-type specificity of significant interactions. (C) Venn diagram illustrating total numbers of interactions called in each cell type. (D) Whole-region view of classified interactions throughout the Sox2 region. (E) Heatmap showing enrichment of selected genomic annotations (rows) within interaction classes (columns) relative to background bin-bin pairs. (F) Enrichment above background of motif orientations of CTCF sites occupied in NPCs found at the base of significant interactions identified in pNPCs.

## Discussion

Recovering bona fide, biologically-relevant looping interactions from 5C data is a challenging problem, requiring multiple steps to account for sequencing depth and library complexity, primer- and fragment-specific biases, spatial noise, distance-dependent background signal, region- and domain-specific contact frequency effects, and statistical variance estimation. Each of these steps can be addressed with any of a wide variety of proposed approaches, the appropriate selection of which can often be critical to the success of the loop identification endeavor. Here, we provide what is, to the best of our knowledge, the first in-depth, systematic overview of available methods for analyzing loops in 5C data. In the process, we introduce several novel approaches and algorithm variants specifically for 5C data, including a variant of conditional quantile normalization, an explicit spline-based normalization method, the Joint Express balancing algorithm, and a quadratic MVR estimation approach.

Many of the decisions made in the 5C analysis process involve critical tradeoffs: increasing noise for increasing resolution, improving condition-specificity for increasing batch effects, and increased precision for potential overfitting. Here, we have highlighted several of the most important tradeoffs, as well as the most important lessons to be learned from a systematic comparison, including the benefits of using a local donut expected model and the importance of over-dispersion in 5C variance modeling. We note that the MVR estimation is not exhaustively discussed and remains an important area for future inquiry. Moreover, the task of classifying differential looping interactions is only discussed briefly here and remains an exciting area for future work. We provide the coding package lib5C to allow investigators to assess the effects of the tradeoffs discussed here on their own novel 5C data sets.

## Acknowledgements

JEPC is a New York Stem Cell Foundation (NYSCF) Robertson Investigator and an Alfred P. Sloan Foundation Fellow. This work was funded by The New York Stem Cell Foundation (J.E.P.C), the Alfred P. Sloan Foundation (J.E.P.C), the NIH Director’s New Innovator Award (1DP2MH11024701; J.E.P.C), a 4D Nucleome Common Fund grant (1U01HL12999801; J.E.P.C) and a joint NSF-NIGMS grant to support research at the interface of the biological and mathematical sciences (1562665; J.E.P.C).

## References

Anders, S., and Huber, W. (2010). Differential expression analysis for sequence count data. Genome Biology 11, 12.

Beagan, J.A., Duong, M.T., Titus, K.R., Zhou, L.D., Cao, Z.D., Ma, J.J., Lachanski, C.V., Gillis, D.R., and Phillips-Cremins, J.E. (2017). YY1 and CTCF orchestrate a 3D chromatin looping switch during early neural lineage commitment. Genome Research 27, 1139–1152.

Beagan, J.A., Gilgenast, T.G., Kim, J., Plona, Z., Norton, H.K., Hu, G., Hsu, S.C., Shields, E.J., Lyu, X.W., Apostolou, E., et al. (2016). Local Genome Topology Can Exhibit an Incompletely Rewired 3D-Folding State during Somatic Cell Reprogramming. Cell Stem Cell 18, 611–624.

Crane, E., Bian, Q., McCord, R.P., Lajoie, B.R., Wheeler, B.S., Ralston, E.J., Uzawa, S., Dekker, J., and Meyer, B.J. (2015). Condensin-driven remodelling of X chromosome topology during dosage compensation. Nature 523, 240–U299.

Daley, T., and Smith, A.D. (2013). Predicting the molecular complexity of sequencing libraries. Nature Methods 10, 325-+.

Dekker, J., Belmont, A.S., Guttman, M., Leshyk, V.O., Lis, J.T., Lomvardas, S., Mirny, L.A., O’Shea, C.C., Park, P.J., Ren, B., et al. (2017). The 4D nucleome project. Nature 549, 219–226.

Deng, L., and Blobel, G.A. (2014). Manipulating nuclear architecture. Current Opinion in Genetics & Development 25, 1–7.

Dixon, J.R., Selvaraj, S., Yue, F., Kim, A., Li, Y., Shen, Y., Hu, M., Liu, J.S., and Ren, B. (2012). Topological domains in mammalian genomes identified by analysis of chromatin interactions. Nature 485, 376–380.

Dostie, J., Richmond, T.A., Arnaout, R.A., Selzer, R.R., Lee, W.L., Honan, T.A., Rubio, E.D., Krumm, A., Lamb, J., Nusbaum, C., et al. (2006). Chromosome Conformation Capture Carbon Copy (5C): A massively parallel solution for mapping interactions between genomic elements. Genome Research 16, 1299–1309.

Hansen, K.D., Irizarry, R.A., and Wu, Z.J. (2012). Removing technical variability in RNA-seq data using conditional quantile normalization. Biostatistics 13, 204–216.

Imakaev, M., Fudenberg, G., McCord, R.P., Naumova, N., Goloborodko, A., Lajoie, B.R., Dekker, J., and Mirny, L.A. (2012). Iterative correction of Hi-C data reveals hallmarks of chromosome organization. Nature Methods 9, 999-+.

Jhunjhunwala, S., van Zelm, M.C., Peak, M.M., and Murre, C. (2009). Chromatin Architecture and the Generation of Antigen Receptor Diversity. Cell 138, 435–448.

Jin, F.L., Li, Y., Dixon, J.R., Selvaraj, S., Ye, Z., Lee, A.Y., Yen, C.A., Schmitt, A.D., Espinoza, C.A., and Ren, B. (2013). A high-resolution map of the three-dimensional chromatin interactome in human cells. Nature 503, 290–294.

Knight, P.A., and Ruiz, D. (2013). A fast algorithm for matrix balancing. Ima Journal of Numerical Analysis 33, 1029–1047.

Langmead, B., Trapnell, C., Pop, M., and Salzberg, S.L. (2009). Ultrafast and memory-efficient alignment of short DNA sequences to the human genome. Genome Biology 10, 10.

Li, G., Fullwood, M.J., Xu, H., Mulawadi, F.H., Velkov, S., Vega, V., Ariyaratne, P.N., Bin Mohamed, Y., Ooi, H.S., Tennakoon, C., et al. (2010). ChIA-PET tool for comprehensive chromatin interaction analysis with paired-end tag sequencing. Genome Biology 11, 13.

Lieberman-Aiden, E., van Berkum, N.L., Williams, L., Imakaev, M., Ragoczy, T., Telling, A., Amit, I., Lajoie, B.R., Sabo, P.J., Dorschner, M.O., et al. (2009). Comprehensive Mapping of Long-Range Interactions Reveals Folding Principles of the Human Genome. Science 326, 289–293.

Marinov, G.K., Kundaje, A., Park, P.J., and Wold, B.J. (2014). Large-Scale Quality Analysis of Published ChIP-seq Data. G3-Genes Genomes Genetics 4, 209–223.

Nora, E.P., Lajoie, B.R., Schulz, E.G., Giorgetti, L., Okamoto, I., Servant, N., Piolot, T., van Berkum, N.L., Meisig, J., Sedat, J., et al. (2012). Spatial partitioning of the regulatory landscape of the X-inactivation centre. Nature 485, 381–385.

Rao, S.S.P., Huntley, M.H., Durand, N.C., Stamenova, E.K., Bochkov, I.D., Robinson, J.T., Sanborn, A.L., Machol, I., Omer, A.D., Lander, E.S., et al. (2014). A 3D Map of the Human Genome at Kilobase Resolution Reveals Principles of Chromatin Looping. Cell 159, 1665–1680.

Rhind, N., and Gilbert, D.M. (2013). DNA Replication Timing. Cold Spring Harbor Perspectives in Biology 5, 26.

Sanyal, A., Lajoie, B.R., Jain, G., and Dekker, J. (2012). The long-range interaction landscape of gene promoters. Nature 489, 109–U127.

Sauria, M.E.G., Phillips-Cremins, J.E., Corces, V.G., and Taylor, J. (2015). HiFive: a tool suite for easy and efficient HiC and 5C data analysis. Genome Biology 16, 10.

Sims, D., Sudbery, I., Ilott, N.E., Heger, A., and Ponting, C.P. (2014). Sequencing depth and coverage: key considerations in genomic analyses. Nature Reviews Genetics 15, 121–132.

Tang, Z.H., Luo, O.J., Li, X.W., Zheng, M.Z., Zhu, J.J., Szalaj, P., Trzaskoma, P., Magalska, A., Wlodarczyk, J., Ruszczycki, B., et al. (2015). CTCF-Mediated Human 3D Genome Architecture Reveals Chromatin Topology for Transcription. Cell 163, 1611–1627.

Yaffe, E., and Tanay, A. (2011). Probabilistic modeling of Hi-C contact maps eliminates systematic biases to characterize global chromosomal architecture. Nature Genetics 43, 1059–U1040.

Zhang, Y., Liu, T., Meyer, C.A., Eeckhoute, J., Johnson, D.S., Bernstein, B.E., Nussbaum, C., Myers, R.M., Brown, M., Li, W., et al. (2008). Model-based Analysis of ChIP-Seq (MACS). Genome Biology 9.

